# Flies Infected by Zombie Fungi in an Urban Fragment

**DOI:** 10.1101/011742

**Authors:** B. C. Barbosa, T. T. Maciel

## Abstract

This study aims to describe associations between the occurrence of entomopathogenic fungi and parasitic flies of the family Muscidae and behaviors presented by these flies parasitized in an urban fragment of semideciduous forest in Brazil. Were recorded two species of entomopathogenic fungi parasitizing flies collected for the first time in the localities, *E. muscae* and O. *sp*. Different from Diptera parasitized by different fungi behaviors were observed. These new records of occurrence, along with data on their hosts, suggest that many species of this group have not yet registered.

## Introduction

The genus *Ophiocordyceps* and *Entomophthora* contains several species of entomopathogenic fungi specialized to infect and kill their hosts due to a development and transmission necessity (Steenberg et al. 2001; Evans et al. 2011). Some species, such as *Ophiocordyceps unilateralis sensu lato*, are capable to manipulate the host’s behavior, causing it to abandon the colony and act in detriment to its own to enhance the parasite’s fitness (Andersen et al. 2009). This behavior, in which genes of the parasite are expressed in the host’s phenotype, may be classified as an extended phenotype (Dawkins 1982).

Due to the high abundance, fungal infections in Formicidae are the most reported and most well known. The infection occurs during foraging when the ants come into contact with the spores that get stuck in their body and penetrate their cuticle. The period of the fungus infection occurs within three to six days after spore adhesion of the body surface ant. Once infected, an individual dies and the fungus produces a fruiting body that grows just behind the head of the parasitized insect. For this structure, the fungus produces spores that are dispersed across the forest floor, where they, infect new hosts (Evans 1982; Evans & Samson 1982, 1984; Pontoppidan et al. 2009).

On the other hand, records the occurrence of infection of entomopathogenic fungi in Diptera are scarce, as well as information about the behavioral changes caused by the fungus and their hosts. Thus, this study describes the occurrence of parasitic associations between entomopathogenic fungi and flies of the family Muscidae and a brief description of their behavior in an urban fragment of Atlantic rainforest in southeastern Brazil.

## Methods and materials

Records occurred in 2014 in Atlantic Rainforest. The first, the Jardim Botânico da Universidade Federal de Juiz de Fora (21 ° 43 ′28 ″S - 43 ° 16′ 47″ W, 750 m a.s.l.), a fragment of Semideciduous Seasonal Forest Montana (Veloso et al. 1991) recently classified by Santiago et al. (2014) as expressive richness, diversity and floristic diversity of woody vegetation, with endangered species with predominance of pioneer plant complex, besides the considerable presence of exotic species. The area of 84 hectares of extension is located within the city limits of Juiz de Fora, southeastern state of Minas Gerais, Brazil.

Fungi were also identified by Prof. Harry Evans, CABI and João Paulo Machado de Araujo of Pennsylvania State University, based on morphological analysis of fruiting bodies and identification of the hosts present.

## Results and discussion

Two species of entomopathogenic fungi were collected for the first time in these localities, *Entomophthora muscae* (Cohn) Fresen. and *Ophiocordyceps sp.* (Berk. & Broome) G.H. Sung, J.M. Sung, Hywel-Jones & Spatafora with host on adults of Diptera (Muscidae).

Due to the effect of partial decomposition caused by the fungus, the identification of Diptera were only possible family level (Muscidae) for the individual parasitized by *Ophiocordyceps sp.* and gender (*Musca* sp) for the individuals parasitized by *Entomophthora muscae*.

In this study, fungal species showed distinct forms as the manipulation of the setting behavior where *E. muscae* seems to control the fly in search of a stem structure, and with his legs cling, favoring for development the fungus. It was observed that the fungus monopolize the entire surface of the fly between the membranes including the abdominal segments, mesossomal and sclerites. In particular, the individual second, was recorded following a major fire in the area after a long period of drought, evidenced the resistances of these fungi in unfavorable environments.

The species *O. sp.* seems manipulate the insect to land on a sheet without the need to hold her where, after the death of the fly, the fruiting body emerges, extrapolated the body of the fly, getting out of the inside through the head.

As data on these parasitic associations are still rare, especially in anthropogenic areas (Travis et al. 1993; Lastra et al. 2006; Sung et al. 2007), these new records of occurrence, along with data on their hosts, suggest that many species of this group have not yet registered. Therefore, more studies in tropical forests will certainly increase the knowledge about these interactions.

